# Epigenetic Modulation of SPCA2 Reverses Epithelial to Mesenchymal Transition in Breast Cancer Cells

**DOI:** 10.1101/2020.11.12.379685

**Authors:** Monish Ram Makena, Myungjun Ko, Donna Kimberly Dang, Rajini Rao

## Abstract

The secretory pathway Ca^2+^-ATPase SPCA2 is a tumor suppressor in triple receptor negative breast cancer (TNBC), a highly aggressive molecular subtype that lacks tailored treatment options. Low expression of SPCA2 in TNBC confers poor survival prognosis in patients. Previous work has established that re-introducing SPCA2 to TNBC cells restores basal Ca^2+^ signaling, represses mesenchymal gene expression, mitigates tumor migration *in vitro* and metastasis *in vivo*. In this study, we examined the effect of histone deacetylase inhibitors (HDACi) in TNBC cell lines. We show that the pan-HDACi vorinostat and the class I HDACi romidepsin induce dose-dependent upregulation of SPCA2 transcript with concurrent downregulation of mesenchymal markers and tumor cell migration characteristic of epithelial phenotype. Silencing SPCA2 abolished the ability of HDACi to reverse epithelial to mesenchymal transition (EMT). Independent of ATPase activity, SPCA2 elevated resting Ca^2+^ levels to activate downstream components of non-canonical Wnt/Ca^2+^ signaling. HDACi treatment led to SPCA2-dependent phosphorylation of CAMKII and β-catenin, turning Wnt signaling off. We conclude that SPCA2 mediates the efficacy of HDACi in reversing EMT in TNBC by a novel mode of non-canonical Wnt/Ca^2+^ signaling. Our findings provide incentive for screening epigenetic modulators that exploit Ca^2+^ signaling pathways to reverse EMT in breast tumors.

**Simple Summary:** The triple receptor negative breast cancer subtype, or TNBC, currently has no tailored treatment options. TNBC is highly metastatic, associated with high patient mortality, and disproportionately occurs in Black/African American women where it contributes to racial disparities in health outcomes. Therefore, we focused on new therapeutic approaches to TNBC. We discovered that levels of the Calcium-ATPase SPCA2 are abnormally low in TNBC and that these low levels correlate with poor survival prognosis in patients. Previously, we showed that recombinant SPCA2 prevented TNBC cells from acquiring aggressive ‘mesenchymal’ properties associated with metastasis both *in vitro* and *in vivo*. These findings motivated us to search for drugs that turn the SPCA2 gene back on in TNBC cells. In this study, we show that histone deacetylase inhibitors increase SPCA2 levels, activate Ca^2+^ signaling and convert cancer cells to a less aggressive ‘epithelial’ state. These findings could lead to new treatment options for TNBC.

**Graphical Abstract:** 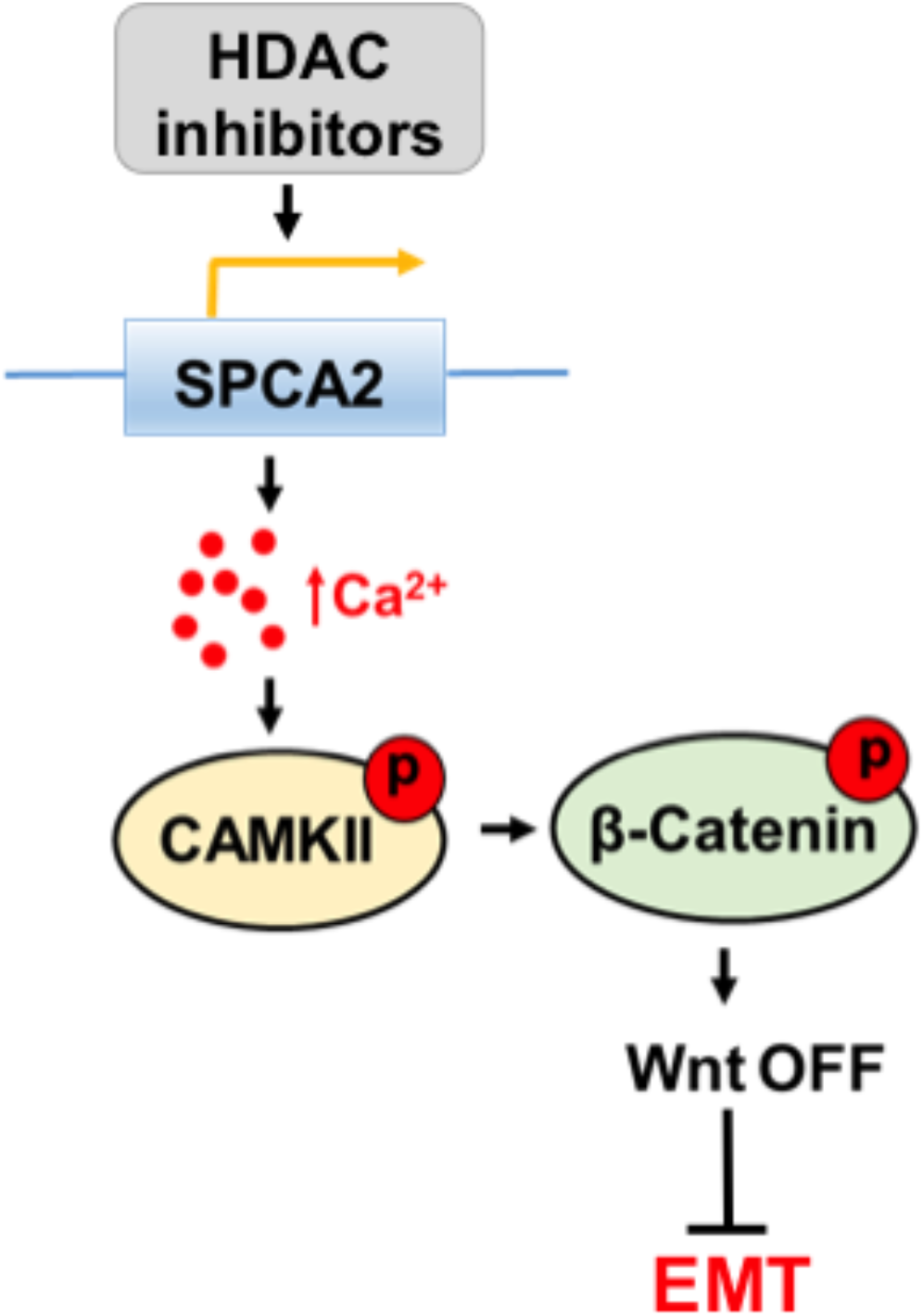

## 1. Introduction

Triple receptor negative breast cancer (TNBC) is a molecular subtype characterized by the lack of estrogen receptor (ER), progesterone receptor (PR) and receptor for human epidermal growth factor-2 (HER2), making it unsuitable for endocrine or targeted antibody therapy [1]. Although TNBC constitutes an estimated 12-15% of all breast cancer subtypes, it is 2.2 times more prevalent in Black/African American women, compared to White/European American women, contributing significantly to the racial disparity in breast cancer outcomes [2]. TNBC tumors are characterized by early relapse and high rates of visceral metastasis [3]. Thus, the lack of tailored treatment options for TNBC, its highly aggressive nature and prevalence in women of color demands urgent focus on new therapeutic approaches.

We recently established a novel role for the secretory pathway Ca^2+^-ATPase SPCA2 as a tumor suppressor in TNBC. Loss of SPCA2 and basal Ca^2+^ signaling in TNBC cells was found to promote epithelial-mesenchymal transition, a hallmark of metastasis, during which cancer cells lose their polarity and cell–cell contacts, thereby acquiring the ability to migrate[4]. At the molecular level, the process of EMT coincides with the loss of epithelial markers, notably the Ca^2+^ binding cell adhesion protein E-cadherin. Conversely, EMT is characterized by gain in expression of mesenchymal genes, including N-cadherin and zinc-finger transcription factors such as Snail, Slug and Zeb1 that reprogram the cancer cell to acquire malignant invasive phenotypes[5]. These gene expression changes serve as convenient markers to track reversible EMT changes in breast cancer cells.

In proof-of-principle experiments we showed that ectopic expression of SPCA2 in TNBC cell lines enhanced post-translational stability of E-cadherin, repressed mesenchymal gene expression and tumor cell migration *in vitro* and mitigated tumor metastasis *in vivo* [4]. Increased surface expression of E-cadherin by SPCA2 activated the tumor suppressor Hippo signaling pathway, resulting in nuclear exclusion and degradation of the transcriptional co-activator YAP (Yes-activated protein) and contact-mediated growth inhibition [4].

Having validated SPCA2 as a potential molecular target in TNBC, our next step is to explore effective treatment options. In this study, we evaluate the efficacy of epigenetic modulators to increase SPCA2 expression in TNBC cell lines. Histone acetyltransferases (HATs) and histone deacetylases (HDACs) counteract each other to regulate gene expression by altering chromatin structure. HDACs remove acetyl groups from lysine residues on histone proteins to condense chromatin and repress gene transcription[6]. Thus, histone deacetylase inhibitors (HDACi) can restore expression of tumor suppressor or endocrine target genes, and a few are being evaluated singly or in combination in clinical trials for the treatment of TNBC/metastatic cancers [7,8]. To date, four HDACi (vorinostat, romidepsin, panobinostat and belinostat) have been approved as second line treatment for relapsed peripheral T-cell lymphoma (PTCL) and/or cutaneous T-cell lymphoma (CTCL) and multiple myeloma by the FDA [6,9]. Alteration in calcium signaling resulted in reactivation of tumor suppressor genes in cancer cells [10]. Some HDACi reportedly regulate intracellular Ca^2+^ levels[11]. At clinically relevant drug concentrations, HDACi have been reported to elevate expression of E-cadherin, while suppressing several mesenchymal genes, thereby decreasing migration *in vitro* and metastasis *in vivo* in triple negative breast cancer cell lines [12–16]. These observations strongly suggest that HDAC inhibitors may upregulate transcription of SPCA2 in TNBC to mediate downstream effects on EMT and metastasis. In this study, we investigated the ability of specific HDAC inhibitors to restore SPCA2 expression and correct dysregulated Ca^2+^ signaling in TNBC.

SPCA2 has dual roles: first, in delivery of Ca^2+^ to the lumen of the Golgi and secretory vesicles where it is important for protein sorting, processing and trafficking, and second, in activating Ca^2+^ entry through Orai1 ion channels for downstream signaling events. ATP hydrolysis is required for Ca^2+^ transport into the Golgi and secretory pathway but not for Orai1 mediated Ca^2+^ entry [17,18]. In this study we distinguish between these dual Ca^2+^ pumping and signaling roles of SPCA2 in mediating mesenchymal to epithelial transition (MET) of TNBC cells. Importantly, we uncover a novel role for SPCA2 in activating downstream components of Wnt/Ca^2+^ signaling pathway. The Wingless-type (Wnt) signaling pathway is an ancient and evolutionarily conserved pathway that regulates embryonic development and drives malignant transformation of cancer cells, leading to EMT, metastasis, and cancer stem cell maintenance [19]. Frequent epigenetic inactivation of Wnt antagonist genes were reported in breast cancer [20]. In the non-canonical Wnt/Ca^2+^ pathway, elevated Ca^2+^ levels activate calcium-binding proteins, including protein kinase C (PKC), calcineurin, and calmodulin-dependent kinase II (CamKII). CaMKII was reported to phosphorylation β-catenin at T332, T472, and S552 [19,21]. Once the β-Catenin is phosphorylated, ubiquitin E3 ligase β-TrCP recognizes phosphorylated β-catenin and promotes its ubiquitination and proteasome degradation, which then promotes MET [22–24]. In this study we demonstrate that upregulation of SPCA2 by HDACi activates downstream components of the Wnt/Ca^2+^ pathway in TNBC cells, which is both necessary and sufficient to mediate key molecular events in MET.

## 2. Results

### 2.1 Low expression of SPCA2 in TNBC correlates with poor survival prognosis

SPCA2 transcript levels are low in triple receptor negative breast cancer derived cell lines[4]. Here, we analyze the TCGA database to show that transcript levels of SPCA2 are significantly (p<0.0001) lower in receptor negative breast tumors compared to receptor positive tumors (Fig. 1A). In contrast, levels of the closely related housekeeping isoform SPCA1 were significantly higher in receptor negative tumors (Fig. 1B). In TNBC, patients with low SPCA2 levels had poorer regression-free survival (Hazard ratio= 0.62 (0.4 – 0.95); Logrank P =0.027; Fig. 1C), whereas this trend is not observed for SPCA1 (Logrank P =0.13; Fig. 1D). These observations suggest that SPCA2 could play a role as a tumor suppressor in TNBC.

**Figure 1:**
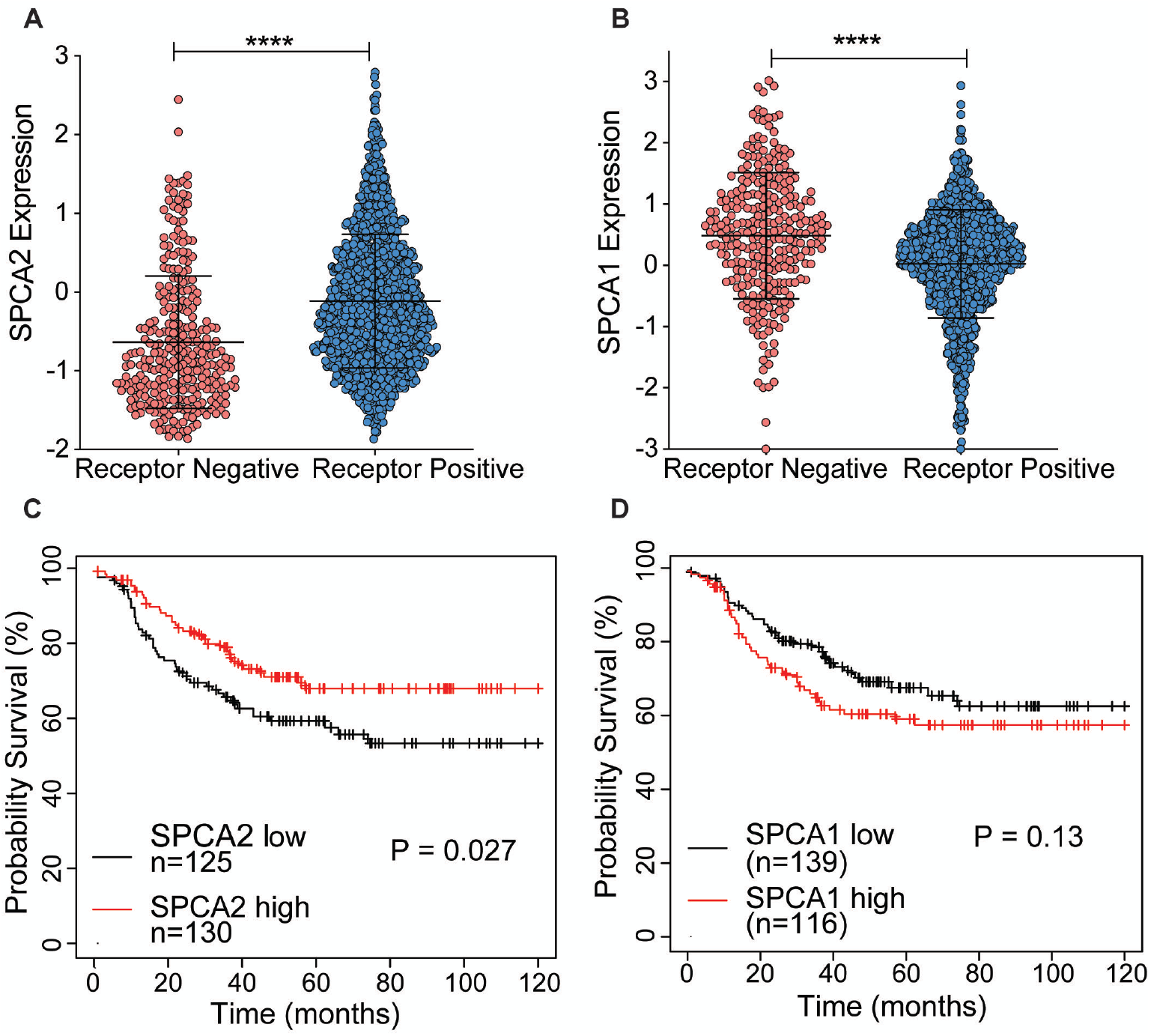
Isoform-specific expression and survival prognosis of SPCA genes in TNBC. (A-B) Expression of (A) SPCA2 and (B) SPCA1 was analyzed in receptor positive and negative (TNBC) breast cancers from TCGA database; n= 290 for receptor-negative patients and n=1222 for receptor-positive patients. (C-D) Kaplan–Meier analysis of regression-free survival in patients stratified by (C) SPCA2 or (D) SPCA1 expression level. Data from KMplotter selected ER, PR, and HER2 (negative). Hazard ratio= 0.62 (0.4 – 0.95); Logrank test, n=125 in low SPCA2 group, and n=130 in high SPCA2 group. Hazard ratio= 1.38 (0.91 – 2.11), Logrank test, n=139 in low SPCA1 group and n=116 in high SPCA1 group.

### 2.2 HDAC inhibitors increase SPCA2 expression in TNBC cells

Previously, we showed that ectopic expression of recombinant SPCA2 (SPCA2R) in TNBC cell lines reduced tumor cell migration *in vitro* and tumor metastasis *in vivo*[4]. These findings prompted a search of pharmacological agents that could increase endogenous SPCA2 gene expression as potential therapeutic approach for TNBC. The addition of acetyl groups to lysine residues on histone proteins modifies chromatin structure and represses gene transcription. Thus, histone deacetylase inhibitors (HDACi) restore expression of tumor suppressor or endocrine target genes [13,25,26] and are being evaluated singly or in combination in clinical trials for the treatment of TNBC/metastatic cancers [8,27,28]. Indeed, epigenetic modulators have been reported to regulate expression of other members of the Ca^2+^-ATPase family[11].

A comparison of histone modifications in the promoter region of the *ATP2C2* gene encoding SPCA2 between normal human mammary epithelial cells (HMEC) and the breast cancer cell line MCF-7 revealed significant elevation of H3K27 histone acetylation in the tumor cells corresponding to high expression of SPCA2 in this cell line[29] (Supplemental Figs. 1A-B). Thus, we posited that HDACi could restore SPCA2 expression in TNBC (Fig. 2A) and evaluated the effect of the pan-HDACi vorinostat and the class I HDACi romidepsin [30] on SPCA2 expression in the metastatic breast cancer cell lines, MDA-MB-231 and Hs578T.

**Figure 2:**
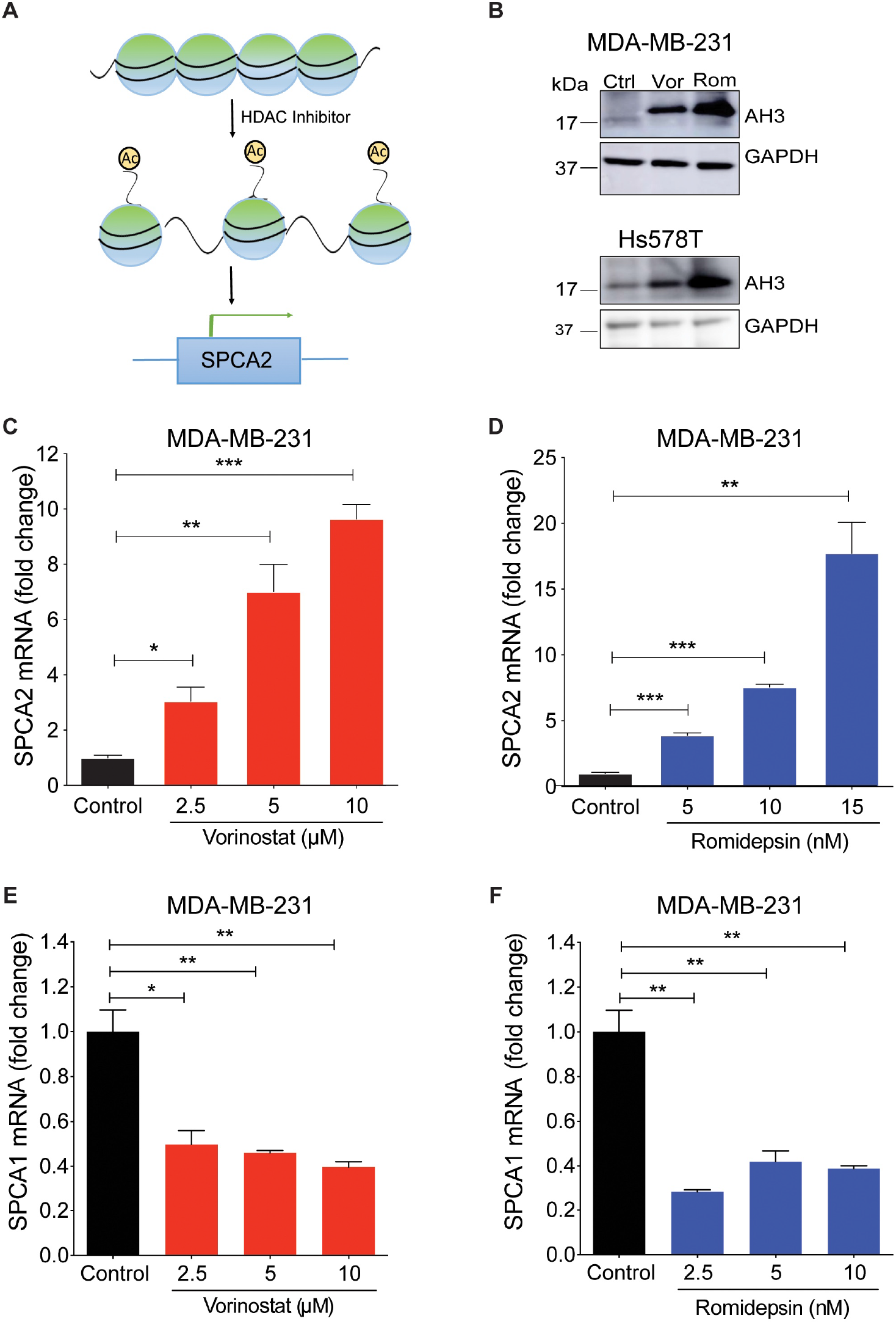
Epigenetic modulation of SPCA2 in TNBC. (A) Schematic showing hypothetical effect of HDAC inhibitors on SPCA2 expression. (B) Histone acetylation in MDA-MB-231 and HS578T cells increased after treatment with vorinostat (2.5 μM) or romidepsin (5 nM) for 24 hours. (C-D) Treatment of MDA-MB-231 cells with HDAC inhibitors vorinostat (2.5, 5, 10 μM) and romidepsin (5, 10, 15 nM) for 24 hours resulted in significant dose-dependence increase of the SPCA2 transcript, n=3. (E-F) Treatment of MDA-MB-231 cells with HDAC inhibitors vorinostat (2.5, 5, 10 μM) and romidepsin (5, 10, 15 nM) for 24 hours resulted in significant decrease of the SPCA1 transcript, n=3. Student’s t-test, ****p<0.0001, ***p<0.001, **p<0.01, *p<0.05.

As expected, vorinostat and romidepsin increased histone acetylation, a hallmark of HDAC inhibitor activity, in both TNBC cell lines (Fig. 2B). We observed a significant dose-dependent up-regulation of SPCA2 transcript with vorinostat and romidepsin with 24 h exposure to clinically relevant drug concentrations in MDA-MB-231 (Figs. 2C and D). SPCA2 transcript increased with longer treatments as seen for romidepsin (5 nM) over 24 and 48 h (Supplemental Fig. 1C). Similar transcriptional changes were observed in Hs578T where both vorinostat (2.5 μM) and romidepsin (5 nM) elicited approximately 5-fold increase in SPCA2 expression (Supplemental Fig. 1D). We found that other classes of epigenetic modulators could enhance SPCA2 expression levels as well. The DNA demethylating agent 5-azacytidine (10 μM) and the natural polyphenol resveratrol (25 μM) which has broad epigenetic regulatory effects, elicited ~3-fold increase of SPCA2 in MDA-MB-231 cells (Supplemental Fig. 1E-F).

In contrast, SPCA1 levels were significantly down regulated by vorinostat and romidepsin in MDA-MB-231 cells (Figs. 2E-F). These opposing transcriptional changes in SPCA1 and SPCA2 illustrate distinct, isoform specific effects of HDACi and are consistent with our previous finding that expression of SPCA1 and SPCA2 is clustered differentially with mesenchymal and epithelial signature gene markers, respectively[4,18].

### 2.3 HDAC inhibitors promote MET transition in TNBC cells

Ectopic expression of SPCA2R was found to diminish mesenchymal gene expression in TNBC cells[4]. Therefore, we evaluated whether concentrations of vorinostat (2.5 μM), shown to induce a modest 3~-fold increase in SPCA2 expression (Fig. 2C), would be sufficient to elicit MET (Fig. 3A). Vimentin is a cytoskeletal intermediate filament protein that is frequently used as a mesenchymal marker. Here we show that treatment of TNBC cell lines with HDAC inhibitors significantly decreased vimentin mRNA (Figs. 3B-C) and protein (Fig. 3D) in MDA-MB-231 and Hs578T cell lines. Decreased cellular expression of vimentin in vorinostat treated MDA-MB-231 cells was confirmed by immunostaining and confocal microscopy (Fig. 3E-F). Vimentin filament assembly in mesenchymal cells is associated with loss of cell contacts and increased motility, which are hallmarks of metastatic progression [31]. We show that treatment of MDA-MB-231 cells with vorinostat significantly decreased migration in Boyden chambers when compared to control (Fig. 3G-H).

**Figure 3:**
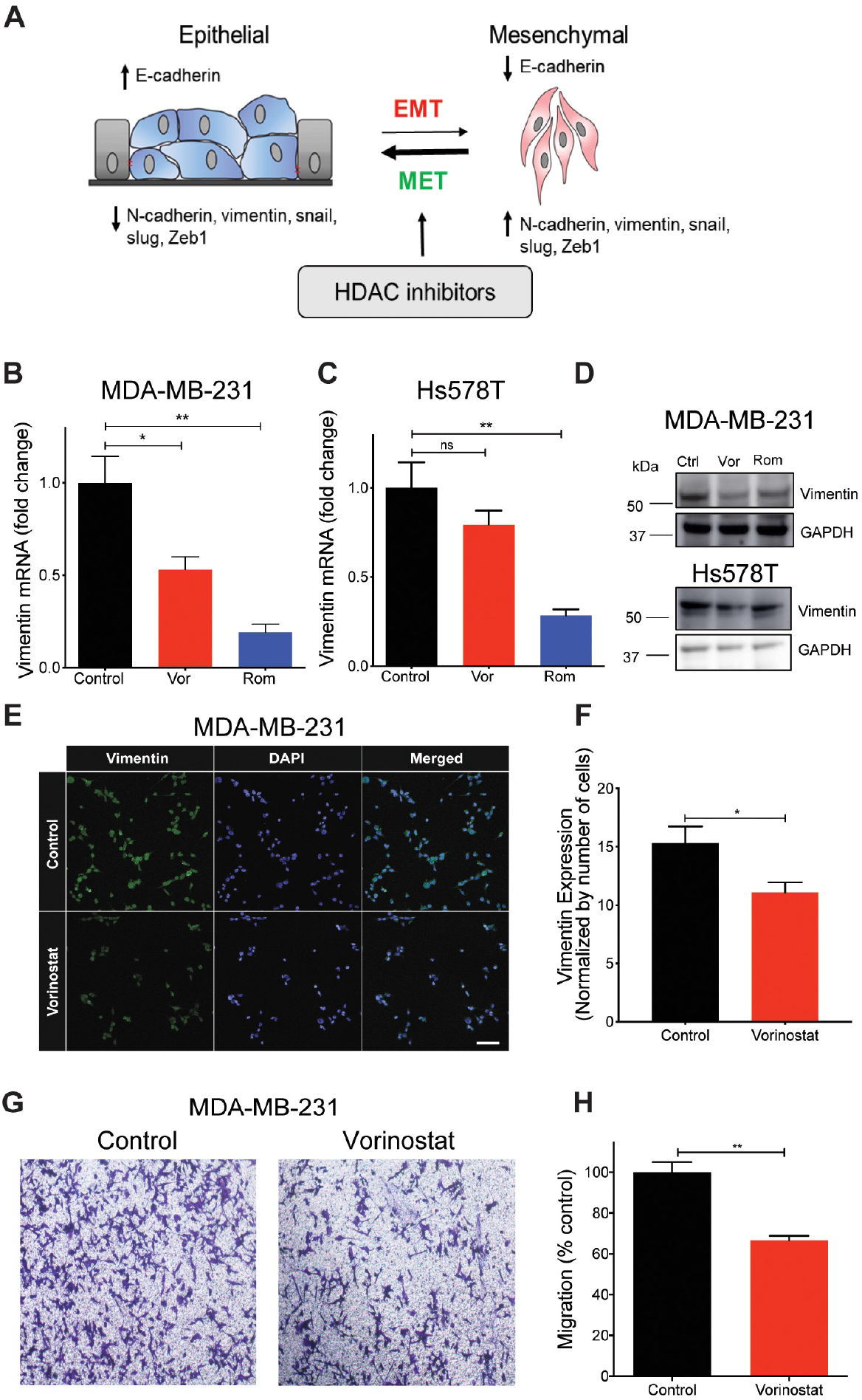
HDAC inhibitors promote MET phenotype TNBC cells. (A) Schematic showing HDAC inhibitors promote MET. Epithelial and mesenchymal states are characterized by gene expression markers that increase or decrease, as indicated, upon transition between states. (B-C) Vimentin mRNA, determined by qPCR, decreased following treatment of MDA-MB-231 (B) and Hs578T (C) with vorinostat (2.5 μM) and romidepsin (5 nM) for 24 hours. n=3. (D) Vimentin protein expression was detected using western blotting in the indicated TNBC cell lines; GAPDH was used as a loading control. (E) Representative confocal microscope images showing immunofluorescence staining of vimentin in MDA-MB-231 cells treated with vehicle (control) or vorinostat (2.5 μM) for 24 hours. (40x magnification; scale bar, 50 μm). (F) Vimentin staining was quantified by ImageJ software, and divided by number (n) of cells, n=352 in control, n= 141 for vorinostat. (G) Representative microscope images of MDA-MB-231 cells treated with vehicle or vorinostat (2.5 μM) for 24 hours, in Boyden chamber (10x magnification). (H) Migration was quantified by ImageJ software from 3 images for each condition. Significance was determined by Student t-test, **p<0.01,*p<0.05, ^ns^ p>0.05.

In addition, HDAC inhibitors suppressed expression of several signature mesenchymal genes, including CDH2 (N-cadherin), SNAI2 (Snail2), and ZEB1 (Zinc Finger E-Box Binding Homeobox 1) in TNBC cell lines (Supplemental Figs. 2A-F). Reciprocally, HDAC inhibitors elevated expression of the epithelial marker E-cadherin (CDH1) in MDA-MB-231, although not in HS578T (Supplemental Figs. 2G and H), where endogenous CDH1 expression is already 7-fold higher compared to MDA-MB-231 (Supplemental Fig. 2I). While these results are consistent with previous findings on HDACi effects[15,16,32], they suggest the intriguing possibility that key therapeutic features associated with HDACi in TNBC cells (Fig. 3A) may be mediated by epigenetic modulation of SPCA2.

### 2.4 SPCA2 is required for HDACi-induced MET changes in TNBC cell lines

Histone deacetylase inhibitors have pleiotropic cellular effects on a wide range of transcriptional and non-transcriptional targets, resulting in alteration of apoptosis, differentiation, signaling and many other pathways central to cancer cell fate and function[33]. We hypothesized that SPCA2 is an upstream target of HDACi that could mediate further downstream effects characteristic of MET (Fig. 4A). To test this hypothesis, we used lentiviral expression of silencing constructs to further reduce the already low levels of SPCA2 transcript in MDA-MB-231 cells[4] (Fig. 4B). SPCA2 knockdown (SPCA2 KD) resulted in the increased expression of mesenchymal genes, including N-cadherin, SNAI1 and vimentin (Supplemental Fig. 3). Similar results were observed earlier in MCF-7 cell line [4]. As expected, vorinostat failed to increase SPCA2 levels in SPCA2 KD cells (Fig. 4C).

Importantly, we show that silencing of SPCA2 prevented downregulation of mesenchymal markers vimentin, N-cadherin and ZEB1 by vorinostat; instead, transcripts of these mesenchymal markers were increased above untreated control (Fig. 4D-F). These reversals were also seen in cellular phenotypes: we show that vorinostat-mediated inhibition of migration was effectively abolished by SPCA2 knockdown (Fig. 4G-H). Further, vorinostat cytotoxicity was significantly abrogated by SPCA2 knockdown (Fig. 4I). These results support our hypothesis (Fig. 4A) of a prominent and novel cellular role for SPCA2 in the efficacy of HDACi in mediating mesenchymal to epithelial transition in TNBC.

**Figure 4:**
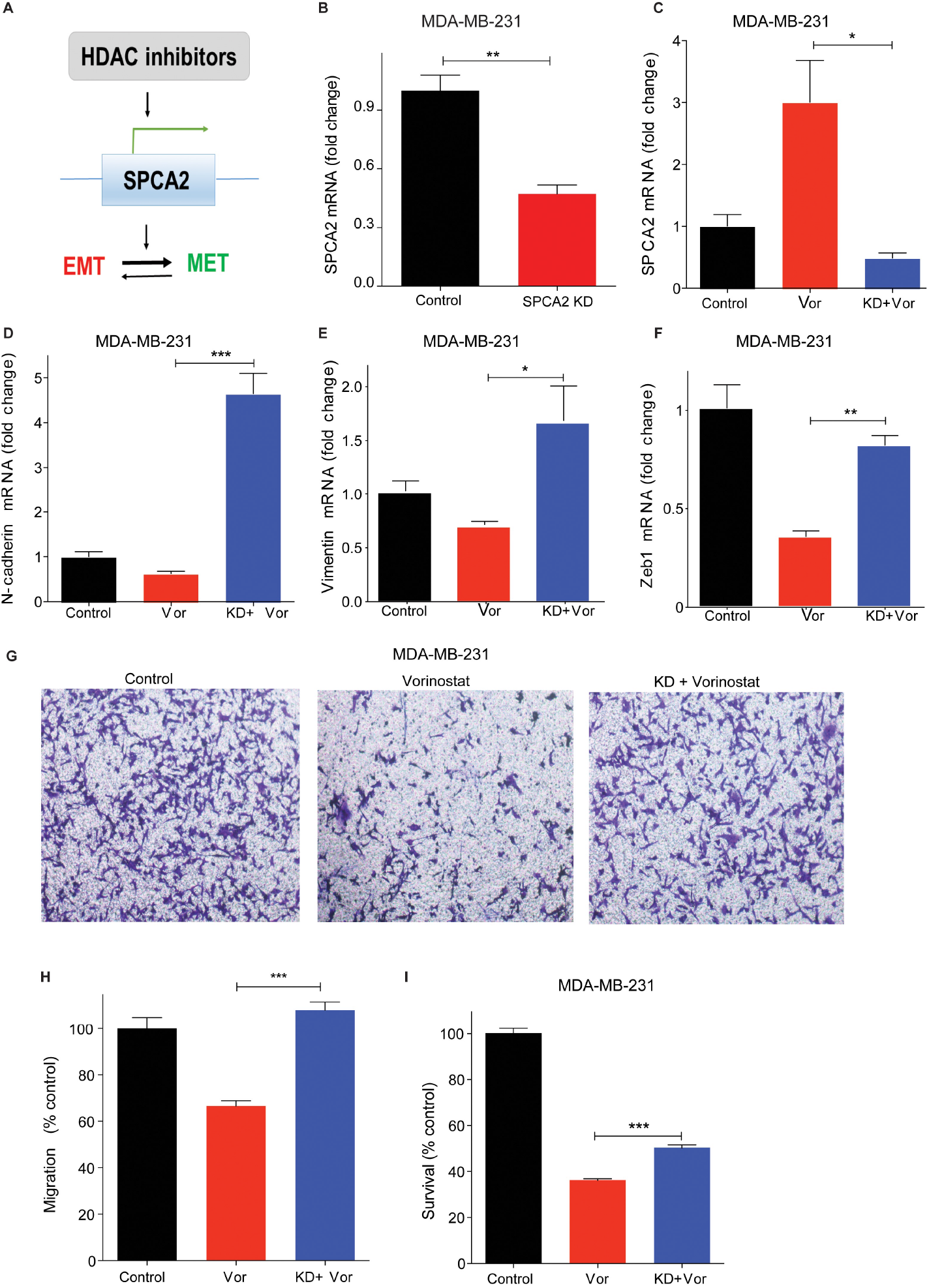
SPCA2 is required for HDACi-induced MET changes in TNBC cell lines. (A) Schematic showing that HDAC inhibitors regulate SPCA2 expression, which in turn, promotes MET. (B) SPCA2 mRNA, determined by qPCR, shows efficacy of knockdown (KD) in MDA-MB-231 cells relative to scramble shRNA control, and (C) following treatment with vehicle (control) or vorinostat (2.5 μM) for 24 hours as indicated. (D-F) Expression of mesenchymal gene markers N-cadherin, vimentin and ZEB1 was determined by qPCR following treatment with vehicle (control) or vorinostat (2.5 μM). SPCA2KD effectively reversed decrease in mesenchymal gene expression in the presence of vorinostat. n=3. (G) Representative microscope images of MDA-MB-231 cells and MDA-MB-231 SPCA2 KD treated with vehicle or vorinostat (2.5 μM) for 24 hours, in Boyden chamber (10x magnification). (H) Migration was quantified by ImageJ software from 3 images for each condition. (I) SPCA2 KD in MDA-MB-231 significantly reduced cytotoxic effect of vorinostat (n=6, 2.5 μM for 80 hours, MTT assay). Student t-test, ***p<0.001, **p<0.01,*p<0.05.

### 2.5 Induction of MET by SPCA2 is independent of Ca^2+^ pumping activity

Similar to other P-type ATPases, the ion pumping activity of SPCA2 requires ATP hydrolysis for the delivery of Ca^2+^ ions into the lumen of the secretory pathway for protein sorting and trafficking [29]. Therefore, we asked if the Ca^2+^-ATPase activity of SPCA2 was required for eliciting MET. A conserved aspartate is essential for formation of the catalytic phosphoenzyme intermediate in all P-type ATPases, including SPCA2. Previously, we showed that mutation of this aspartate in SPCA2 (D379N) inactivated Ca^2+^ pumping activity without affecting protein expression and Golgi localization [29,34]. Here we show that ectopic expression of SPCA2 mutant D379N [34], suppresses mesenchymal gene markers, including vimentin (Fig. 5A-B), N-cadherin, SNAI1 and SNAI2, and ZEB1 (Fig. 5C-F), similar to wild type SPCA2R, reported previously[4].

**Figure 5:**
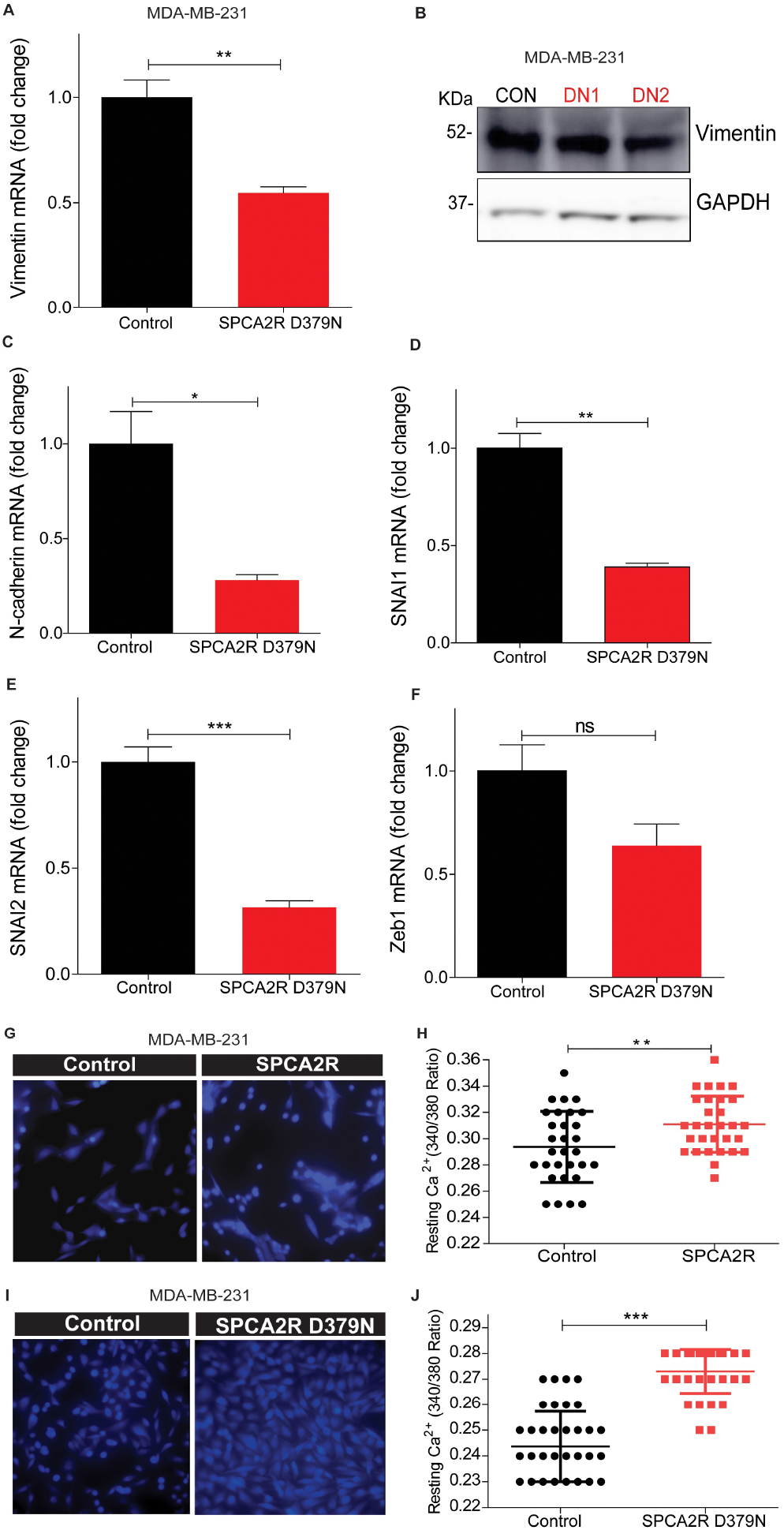
Induction of MET by SPCA2 is independent of Ca^2+^-ATPase activity. (A-B) Ectopic expression of SPCA2 mutant D379N in MDA-MB-231 significantly decreased vimentin compared to control, as determined by (A) qPCR (n=3) or (B) Western blotting. GAPDH was used as a loading control. (C-F) Mesenchymal gene markers, measured by qPCR, were decreased following ectopic expression of SPCA2R D379N as shown. n=3. (G-J) Representative live cell Ca^2+^ imaging baseline readings in Ca^2+^-free conditions using Fura2-AM treated MDA-MB-231 cells with or without SPCA2R (G) or D379N (I). Fluorescence emission ratio of 340/380 nm showing increase in average resting Ca^2+^ in the presence of (H) SPCA2R (control n=30 cells, SPCA2R n=30 cells) or (J) D379N (control n=40 cells, SPCA2R D379N n=40 cells). Student t-test; ***p<0.001, **p<0.01,*p<0.05, ^ns^ p>0.05.

Unlike other P-type ATPases, SPCA2 has a unique function in mediating store independent Ca^2+^ entry (SICE)[29,35], so called because it is distinct from store operated Ca^2+^ entry (SOCE) that occurs upon depletion of endoplasmic reticulum Ca^2+^ stores. Independent of ATPase activity, SPCA2 chaperones and activates the Orai1 Ca^2+^ channel to elevate resting Ca^2+^ levels in the cytoplasm[29]. We ectopically expressed recombinant SPCA2 (SPCA2R) or the catalytically inactive D379N mutant in MDA-MB-231 cells and imaged live cells using the ratiometric Ca^2+^ indicator Fura-2 (Fig. 5G, I). As seen by the emission ratio from 340 and 380 nm excitation (Fig. 5H, J), D379N elicited the elevation of resting cytoplasmic Ca^2+^ similar to SPCA2R control. These findings suggest that SPCA2 mediates MET through an elevation of cytoplasmic Ca^2+^.

### 2.6 SPCA2 activates the Wnt/Ca^2+^ signaling pathway in TNBC cells

Wnt signaling is a well-known regulator of cell fate, including self-renewal, proliferation, differentiation and apoptosis, both in normal mammary development and in breast cancer[36]. The key effector protein in canonical Wnt signaling is β-catenin. When the Wnt pathway is OFF, β-catenin is phosphorylated and degraded. When Wnt is ON, β-catenin can translocate to the nucleus to activate genes that promote self-renewal and stemness associated with the mesenchymal state. In the non-canonical Wnt/Ca^2+^ pathway, a rise in cytosolic Ca^2+^ has an antagonistic effect on canonical Wnt signaling. Wnt-mediated release of Ca^2+^ from endoplasmic reticulum stores phosphorylates Ca^2+^-calmodulin activated kinase, CAM Kinase II, leading to phosphorylation and degradation of β-catenin, turning Wnt signaling OFF [37]. We hypothesized that elevation of basal Ca^2+^ by SPCA2 could directly activate downstream components of Wnt/Ca^2+^ signaling to mediate MET (Fig. 6A). Ectopic expression of SPCA2R in MDA-MB-231 increased phosphorylation of CAM Kinase II, by 2-fold, and increased phosphorylation of β-catenin by 2.5-fold (Fig. 6B-D). Consistent with these findings, addition of 2 mM extracellular Ca^2+^ to cells maintained in nominally Ca^2+^-free culture medium increased phospho-CAMK II and decreased non-phospho (active) β-catenin and total β-catenin levels revealing a novel role for extracellular Ca^2+^ in Wnt/Ca^2+^ signaling (Supplementary Fig. 4). To determine if SPCA2R-mediated CAMK II signaling was important for MET related gene expression changes, we treated cells with the CAMK II inhibitor CKI. We found that transcriptional downregulation of N-cadherin, vimentin and ZEB1 by SPCA2R expression in MDA-MB-231 cells was reversed by CKI treatment (Fig. 6E-G).

**Figure 6:**
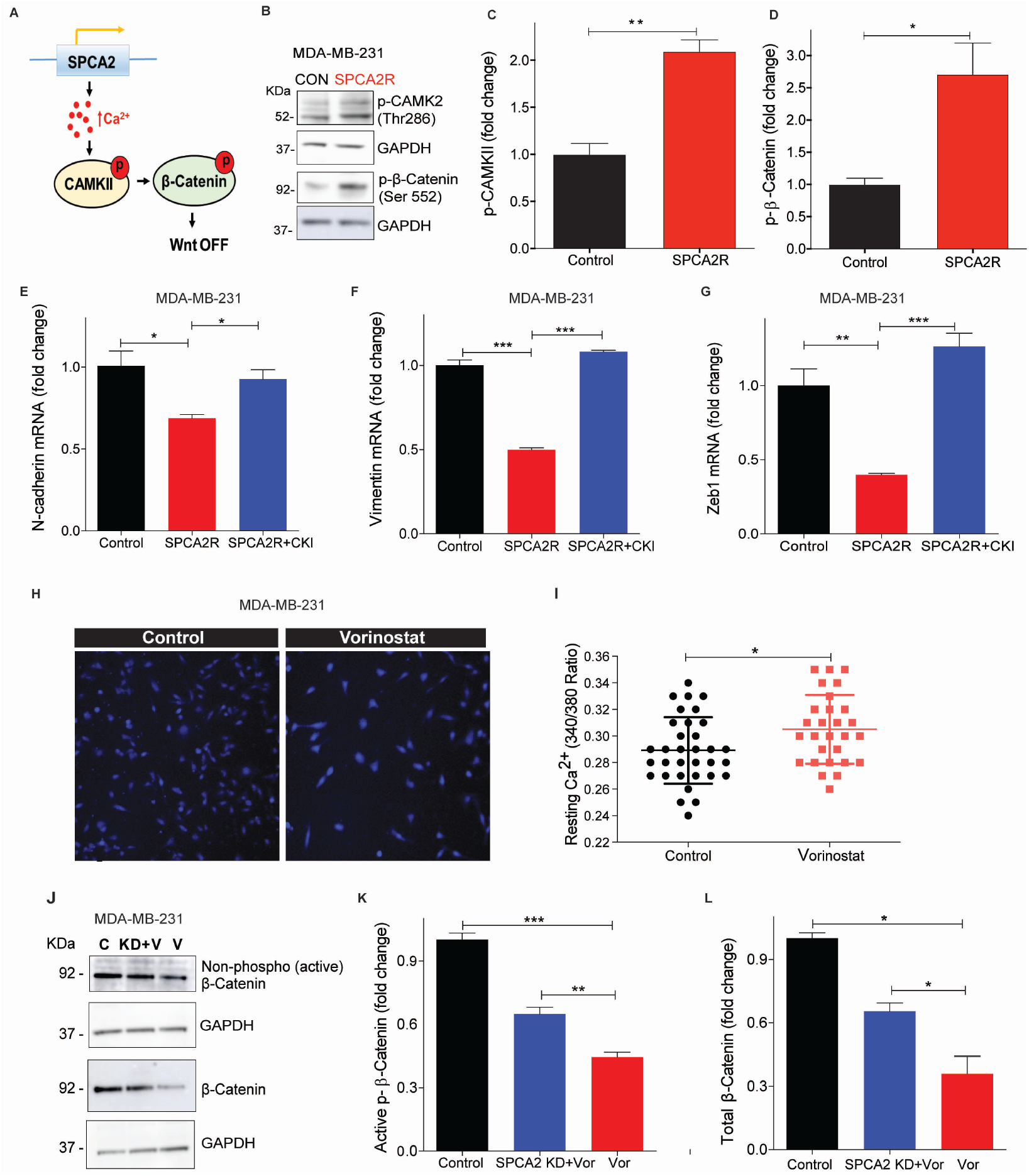
SPCA2 activates downstream Wnt/Ca^2+^ signaling. (A) Schematic of downstream activation of Wnt/Ca^2+^ signaling by SPCA2. Store-independent Ca^2+^ entry activated by SPCA2 increases cytoplasmic Ca^2+^, activating CAMKII and inactivating β-catenin by phosphorylation, turning Wnt pathway OFF. (B) Representative Western blot of p-CAMKII and p-β-catenin in MDA-MD-231 transfected with vector control or SPCA2R; GAPDH was used as a loading control. (C-D) Quantitation of Western blots (n=3) by densitometry. (E-G) Ectopic expression of SPCA2 (SPCA2R) in MDA-MB-231 decreased the mesenchymal gene markers, N-cadherin, vimentin and Zeb1, which were significantly reversed by addition of CAM Kinase II inhibitor (KN 93 phosphate, 20 μM, 24 hours). n=3. (H) Representative live cell Ca^2+^ imaging in calcium-free conditions using Fura2-AM in MDA-MB-231 treated with vehicle (control) or vorinostat (2.5 μM for 24 hours). (I) Fluorescence emission ratio of excitation at 340/380 nm showing increase in average resting Ca^2+^ in the presence of vorinostat (n=28 cells) compared to vehicle control (n=33 cells). (J) Representative Western blot of non-phosphorylated (active) and total β-catenin in MDA-MD-231 control (scrambled shRNA) and SPCA2 KD, treated with vorinostat as indicated; GAPDH was used as a loading control. (K-L) Quantitation of Western blots by densitometry (n=3). SPCA2 KD significantly reversed the decrease in active and total levels of β-catenin by vorinostat. Student t-test, ***p<0.001, **p<0.01,*p<0.05.

Next, we asked if HDACi-mediated increase in SPCA2 expression was sufficient to increase resting Ca^2+^ levels. We show that treatment of MDA-MB-231 with vorinostat (2.5 μM) resulted in increased resting Ca^2+^ levels similar to ectopic SPCA2R expression (Fig. 6H-I). Vorinostat decreased levels of active, non-phosphorylated β-catenin as well as total β-catenin levels (Fig. 6J-L), suggesting that Wnt signaling was turned OFF. However, knockdown of SPCA2 impaired the ability of vorinostat to decrease active and total β-catenin (Fig. 6J-L). Taken together, our results reveal an exciting new role for SPCA2 as an activator of Wnt/Ca^2+^ signaling in TNBC.

## 3. Discussion

Histone modification by acetylation is enriched in transcriptionally active promoter and enhancer regions of the genome where it promotes decondensation of chromatin and facilitates access and recruitment of transcription factors [38]. HDAC inhibitors that enhance acetylation and gene transcription are currently being evaluated in various stages of clinical trials for TNBC/metastatic cancers (NCT02890069, NCT02708680, NCT04315233 and NCT04296942) (https://clinicaltrials.gov). Although the FDA has approved the use of HDACi to treat some aggressive cancers, many patients eventually relapse after treatment. Furthermore, the efficacy of HDAC inhibitors in solid tumors is limited[39]. We have previously established a novel role for the secretory pathway Ca^2+^-ATPase SPCA2 as a tumor suppressor in highly metastatic TNBC cells[4]. This study demonstrates that TNBC tumors with low SPCA2 expression benefit from HDACi treatment that transcriptionally elevates SPCA2 and restores deficient Ca^2+^ signaling to promote MET phenotypes *in vitro*. Thus, stratification of TNBC by SPCA2 expression could be useful in predicting the clinical efficacy of HDACi treatment.

Frequent epigenetic inactivation of Wnt antagonist genes were reported in breast cancer which suggest that their loss of function contributes to activation of Wnt signaling in breast carcinogenesis[20]. For example, APCL, homologous to the adenomatous polyposis coli (APC) tumor suppressor gene, is known to regulate Wnt/β-catenin pathway by promoting β-catenin ubiquitylation and degradation. HDAC inhibitors inhibited tumor growth and metastasis via upregulating APCL expression in breast cancer cells[40]. HDAC inhibitors reversed EMT by inhibiting Epithelial Cell Adhesion Molecule (EpCAM) cleavage and WNT signaling in breast cancer cells[12]. When combined with Wnt/β-catenin pathway inhibitors, HDACi caused a dramatic decrease in cell viability by inducing the extrinsic apoptotic pathway in lung carcinoma cells[41]. In colon cancer cells, HDAC inhibitors were reported to induce cell cycle arrest and apoptosis by upregulating non-canonical Wnt signaling [42]. These results highlight that that HDAC inhibitors induce cell death and inhibit EMT through Wnt/β-catenin pathway.

The endoplasmic reticulum acts as a store of intracellular calcium that can be rapidly released into the cytoplasm to trigger a variety of cellular responses [43]. The Wnt/Ca^2+^ pathway emerged with the finding that some Wnts and Fz receptors can stimulate intracellular Ca^2+^ release from ER, and this pathway is dependent on G-proteins[44,45]. Here, we identify a novel mechanism of activating Wnt/Ca^2+^ signaling by extracellular Ca^2+^ and SPCA2-mediated store independent Ca^2+^ entry (SICE). The resulting elevation of cytoplasmic Ca^2+^ activates CAMK II signaling, leading to phosphorylation of β-catenin, which targets it for inactivation and degradation. We showed that SPCA2-mediated activation of the downstream components of Wnt/Ca^2+^ signaling was necessary and sufficient to promote hallmarks of MET in TNBC cells. It is worth noting that this newly identified pathway may be Wnt ligand independent or there may be other ways to activate SICE that remain to be determined.

Our findings, summarized in the emerging model shown in Fig. 7, provide novel mechanistic insights on druggable pathways that could be harnessed for treatment of metastatic cancers. We have previously shown that ectopic expression of SPCA2 in triple-negative breast cancer elevates baseline Ca^2+^ via SICE leading to phosphorylation and nuclear exclusion of YAP and suppression of EMT[4]. Multiple reports have shown YAP/β-catenin signaling axis as key regulator in cancer progression in variety of cancers [46–50]. Therefore, SPCA2-YAP-β-catenin pathway warrants further investigation.

**Figure 7:**
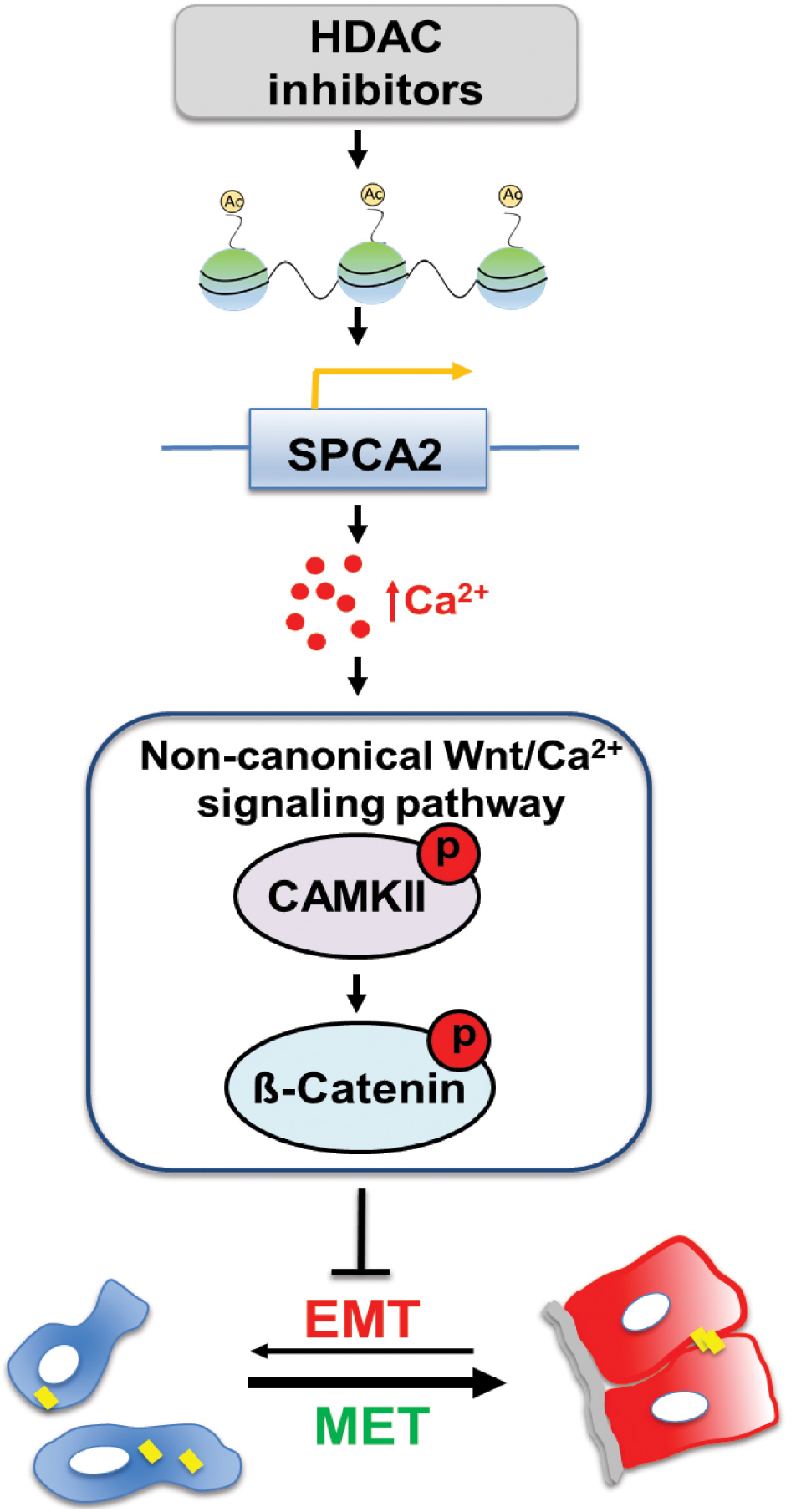
Epigenetic modulation of SPCA2 in TNBC reverses EMT through Ca^2+^/Wnt signaling. Breast cancer cells undergo reversible transitions between mesenchymal state, characterized by low cytosolic Ca^2+^ (blue) and detached junctional protein E-cadherin (yellow bars), and epithelial state, characterized by higher resting Ca^2+^ (red), attachment to the basement membrane (gray) and cell-cell junctions (yellow). Previous work showed that ectopic expression of SPCA2 in TNBC cells increases cytosolic Ca^2+^ and E-cadherin expression to promote epithelial phenotypes. In this study, we show that HDAC inhibitors (vorinostat and romidepsin) increase histone acetylation leading to increased SPCA2 transcription, which in turn elevates resting Ca^2+^ by activation of store-independent Ca^2+^ entry. Ca^2+^ activates non-canonical Wnt signaling by phosphorylating CAMKII and β-catenin, turning Wnt OFF. This blocks EMT and promotes MET.

## 4. Materials and Methods

### 4.1 Chemicals

KN-93 phosphate (CAMK II inhibitor) was obtained from Tocris Bioscience (Bristol, United Kingdom), 5-Azacytidine from Selleck chemicals (Houston, TX), and resveratrol from Sigma Aldrich (St. Louis, MO).

### 4.2 Media and Cells

#### 4.2.1 Cell Culture

Cells used in this study were MDA-MB-231 (ATCC HTB-26) and Hs578T (ATCC HTB-126). MDA-MB-231 and Hs578T cells were gown in DMEM (Thermo Fisher Scientific, Waltham, MA) containing antibiotic, and 5-10% FBS (Sigma Aldrich). Cells were cultured with 5% CO_2_ and at 37°C in a humidified incubator. All cell lines were grown for less than a month or no more than 6 passages. We did not see visible contamination by mycoplasma. Since SPCA2 expression is regulated by cell density, cells were maintained at 50-60% confluency in both knockdown experiments and controls. Although we used lentiviral transfection, resistant clones tended to appear after 2-3 passages, therefore all experiments used early passage cells.

#### 4.2.2 MTT assay

Cells were seeded at ~ 3000 cells/well in 96 well plates. After the cells were incubated overnight, they were treated with vorinostat for 80 hours. Thereafter, MTT dye (Thermo Fisher Scientific) was added to the cells for four hours, and SDS-HCl solution was added to each well and mixed thoroughly. After four hours of incubation, absorbance was measured at 570 nm. Cell survival is calculated as the percentage normalized to control.

### 4.3 HDAC inhibitor treatment

Vorinostat (Vor) was obtained from cell signaling technology (Danvers, MA) and romidepsin (Rom) from Selleck Chemicals. Stock solutions of vorinostat and romidepsin were prepared in DMSO. Cell lines were seeded into 6-well plates and incubated overnight, cells were treated with HDAC inhibitors when they reached ~30-40% of confluency. For dose-dependent experiments, cells were treated with 2.5 μM, 5 μM and 10 μM of vorinostat, and 5 nM, 10 nM, and 15 nM of romidepsin. To investigate changes in EMT markers, MDA-MB-231 and HS578T cells were treated 2.5 μM of vorinostat and 5 nM of romidepsin. Following 24 hours of incubation, RNA was collected and qPCR was performed. Three replicate wells were tested per concentration.

### 4.4 Epigenetic modifications in ATP2C2 promoter region

Epigenetic modifications in the promoter of SPCA2 were obtained from the UCSC Genome Browser Database (http://genome.ucsc.edu/) and Eukaryotic Promoter Database (https://epd.epfl.ch//index.php). Histone methylation and acetylation data was sourced from ENCODE through the IGV browser (https://igv.org/app/).

### 4.5 Molecular Biology Techniques

#### 4.5.1 Lentiviral Transfection

FUGW overexpression constructs, pLK0.1 shRNA lentiviral construction of SPCA2 was packaged and transfected according to previous methods[4], using pCMV-Δ8.9 and PMDG in HEK293T cells. Virus was collected after 24 hours for 3 consecutive days and concentrated with Lenti-X Concentrator (Takara Bio USA, Mountain View, CA). A mixture of two shRNA constructs for SPCA2 was used. Cells were transfected with virus for 48 hours, and selected with puromycin (1 μg/mL for MDA-MB-231). Since the basal SPCA2 expression is low in MDA-MB-231 cell line, the cells were transfected with knockdown virus 2-3 times to get a good efficiency. Knockdowns were confirmed by qPCR. Experiments were performed within 2-3 passages to ensure that knockdown was maintained.

#### 4.5.2 cDNA synthesis and quantitative PCR

1μg of RNA was collected and used for cDNA synthesis (Applied Biosystems, Foster City, CA). The qPCR mastermix was obtained from Thermo Fischer Scientific, 50ng of cDNA, and Taqman probe (Thermo Fischer Scientific) as specified; GAPDH (Hs02796624_g1), SPCA2 (Hs00939492_m1), SPCA1 (Hs00995930_m1), CDH1 (HS01023895_m1), CDH2 (Hs00983056_m1), ZEB1 (Hs00232783_m1), SNAI2 (Hs00161904_m1), and Vimentin (Hs00958111_m1).

### 4.6 Western blotting

Cells were collected, and lysed in RIPA buffer (Thermo Fisher Scientific) supplemented with protease inhibitor cocktail, and phosphatase inhibitors. Protein quantification was done by the bicinchoninic acid assay. Thirty μg of protein in each sample was resolved by electrophoresis using 4-12% bis-tris gels (Thermo Fisher Scientific) and transferred to nitrocellulose membrane (Bio-Rad, Hercules, CA). Immunoblots were probed with antibodies (1:1000) from Cell Signaling Technologies, followed by incubation with HRP-conjugated secondary antibodies. Proteins were visualized using chemiluminescence substrate. Blots were analyzed using ImageJ.

### 4.7 Imaging

#### 4.7.1 Immunofluorescence

Cells were cultured on glass coverslips and were rinsed with PBS and pre-extracted with 1X PHEM buffer, 8% sucrose and 0.025% saponin. Cells were fixed with 4% paraformaldehyde for 30 minutes and were rinsed and washed with PBS 3 times for 5 minutes each. After blocking in 1% BSA, cells were incubated with overnight in 4°C using vimentin antibody (Cell Signaling Technologies). Cells were rinsed with 0.2% BSA 3 times for 5 minutes and were then incubated with a fluorescent secondary antibody in 1% BSA and 0.025% saponin buffer for 30 minutes at room temperature. Coverslips were washed and mounted onto slides with mounting media (Agilent, Santa Clara, CA). Immunofluorescent staining was analyzed using ImageJ.

#### 4.7.2 Live cell calcium imaging

Live imaging of Ca^2+^ was performed using Fura2-AM (Invitrogen, Carlsbad, CA). MDA-MB-231 cells were transfected with vector (Control) or SPCA2R or were treated with vorinostat for 24 hours, washed and treated with Fura2-AM in imaging buffer (20 mM Hepes, 126 mM, NaCl, 4.5 mM KCl, 2 mM MgCl_2_, 10 mM glucose at pH7.4) for 30 min. Cells were excited at 340 nm and 380 nm, and Fura emission was captured at 505 nm [4].

### 4.8 Boyden chamber assay

MDA-MB-231 cells were maintained in DMEM media without FBS for 4 hours. Approximately 5×10^4^ cells of each condition were plated in 100 μL DMEM media without FBS in 6.5 mm Transwell chambers with 8.0 mm Pore Polyester Membrane Insert (Corning Inc., Corning, NY), and transferred to a 24-well dish containing 500 μL fresh media (0.5% FBS containing DMEM) with/without vorinostat. After 24 hours, cells were fixed with 4% paraformaldehyde for 30 minutes, then 0.5% crystal violet (Sigma-Aldrich) was added for 1 hour, and washed with PBS. Using a microscope (Nikon, Eclipse TS100), photos of membrane insert were captured at 10X magnification. ImageJ software was used to quantify migration.

## 5. Conclusions

Ionized Ca^2+^ is a versatile, essential and ubiquitous second messenger that impacts all aspects of cell fate and function, including key hallmarks of cancer such as proliferation, differentiation, migration, invasion, and apoptosis. With the advent of pharmacogenomics and molecular profiling, tailored tumor treatment is now possible with a potential for new and improved therapies aimed at Ca^2+^ signaling pathways and genes. Armed with a recent mechanistic understanding of the subtype-specific role of SPCA2 in Ca^2+^ signaling [18], we explored innovative therapeutic regimens for metastatic breast cancer. We showed that FDA-approved HDACi, vorinostat and romidepsin, were effective in increasing SPCA2 levels at clinically relevant concentrations and that SPCA2 induction was necessary for reversal of EMT phenotypes through downstream Wnt/Ca^2+^ signaling. Although we focused on HDAC inhibitors in this study, we show that other classes of epigenetic modulators, such as DNA methyltransferase inhibitors, could also be effective in upregulating SPCA2, making us optimistic that this approach will be broadly useful. Alternatively, drugs that bypass the role of SPCA2 to directly activate Ca^2+^ channels could also restore basal Ca^2+^ signaling and reverse EMT in TNBC cells. Our findings provide incentive for screening of new or repurposed drugs that exploit subtype-selective Ca^2+^ signaling pathways to reverse EMT in breast tumors.

## Supplementary Materials

The following are available online at www.mdpi.com/xxx/s1, Figure S1: Epigenetic modification in SPCA promoter region; Figure S2: HDAC inhibitors promote MET phenotype TNBC cells; Figure S3: SPCA2 KD induced EMT changes in TNBC cell lines; Figure S4: Extracellular Ca^2+^ entry activates non-canonical Wnt/Ca^2+^ pathway; Figure S5: Uncropped western blots for Figures 2, 3 and 5; Figure S6: Uncropped western blots for Figure 6.

## Author Contributions

Conceptualization, M.R.M. and R.R.; methodology, M.R.M, M.J.K, D.K.D and R.R; software, M.R.M and RR; validation, M.R.M, M.J.K, D.K.D and R.R.; formal analysis, M.R.M, M.J.K, D.K.D and R.R.; investigation, M.R.M, M.J.K, D.K.D and R.R.; resources, R.R.; data curation, M.R.M, M.J.K, D.K.D and R.R.; writing and editing, M.R.M and R.R.; visualization, M.R.M and RR; supervision, R.R.; project administration, R.R.; funding acquisition, M.R.M. and R.R. All authors have read and agreed to the published version of the manuscript.

## Acknowledgments

M.R.M. acknowledges the support of American Association of Cancer Research (AACR). M.J.K acknowledges the support of the graduate training programs in Cellular & Molecular Medicine, at the Johns Hopkins University. Authors like to thank Dr. Vijender Singh, Associate Director of Computational Biology Core, University of Connecticut, for his suggestions regarding epigenetic modification of promoter region.

