## Supplemental Figures for "Epigenetic Modulation of SPCA2 Reverses Epithelial to Mesenchymal Transition in Breast Cancer Cells"

**SUPPLEMENTAL DATA**

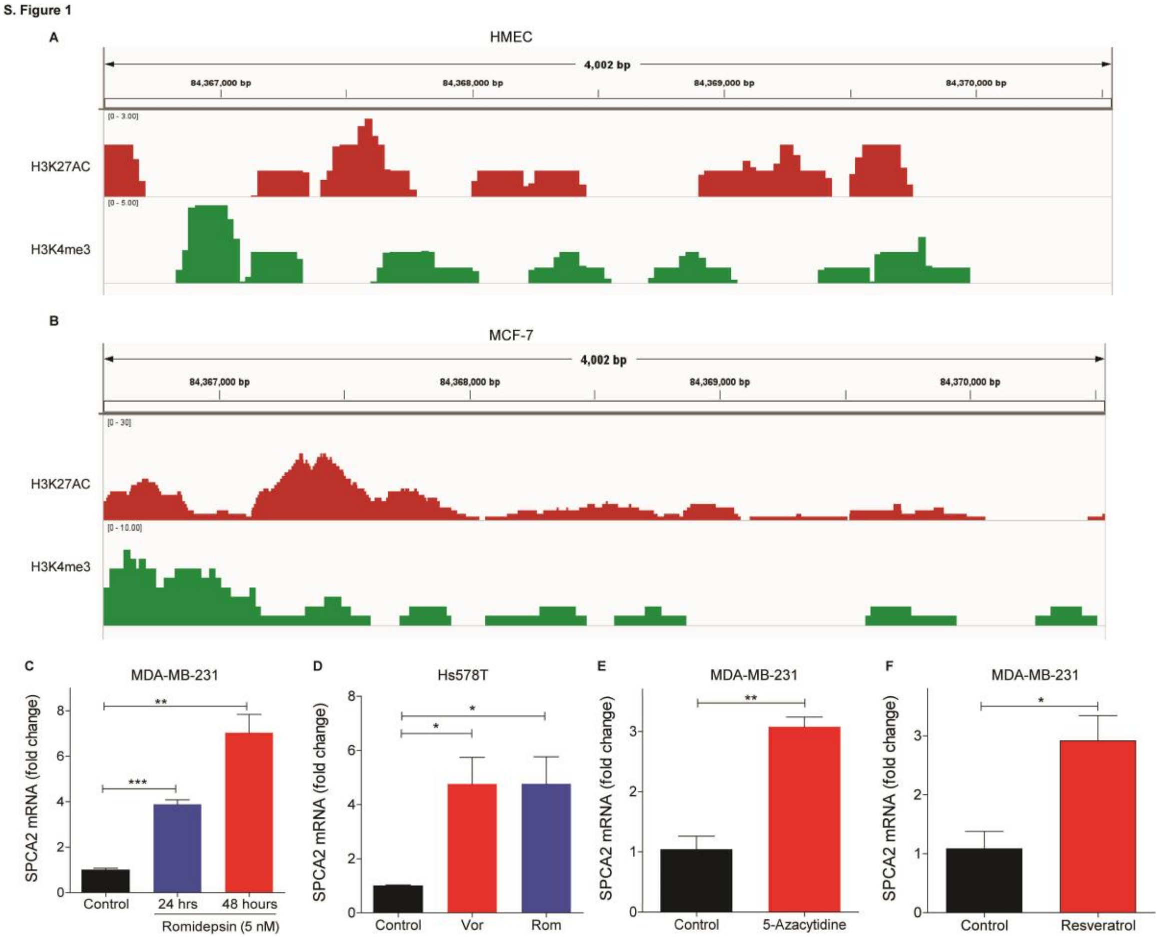


**Supplementary Figure 1: Epigenetic modification in SPCA promoter region.**
(A-B) Epigenetic modification of the promoter region of SPCA2, showing histone acetylation (H3K27AC) and methylation (H3K4me3), in (A) HMEC and (B) MCF-7 cells, as described in Methods. Not the scale bar differences, showing increased acetylation and methylation in MCF-7, compared to HMEC. (C) Treatment of MDA-MB-231 cells with HDAC inhibitors romidepsin showed a time-dependent increase in SPCA2 expression. n=3. (D) Treatment of HS578T cells with HDAC inhibitors vorinostat (2.5 μM) and romidepsin (5 nM) for 24 hours significantly increased SPCA2. (E-F) Treatment of MDA-MB-231 cells with DNA methyltransferase inhibitor (5-Azacytidine, 10 μM, 48 hours) and sirtuin activator (Resveratrol, 2.5 μM, 24 hours) significantly increased SPCA2 n=3. Student's t-test, ***p<0.001, **p<0.01,*p<0.05.

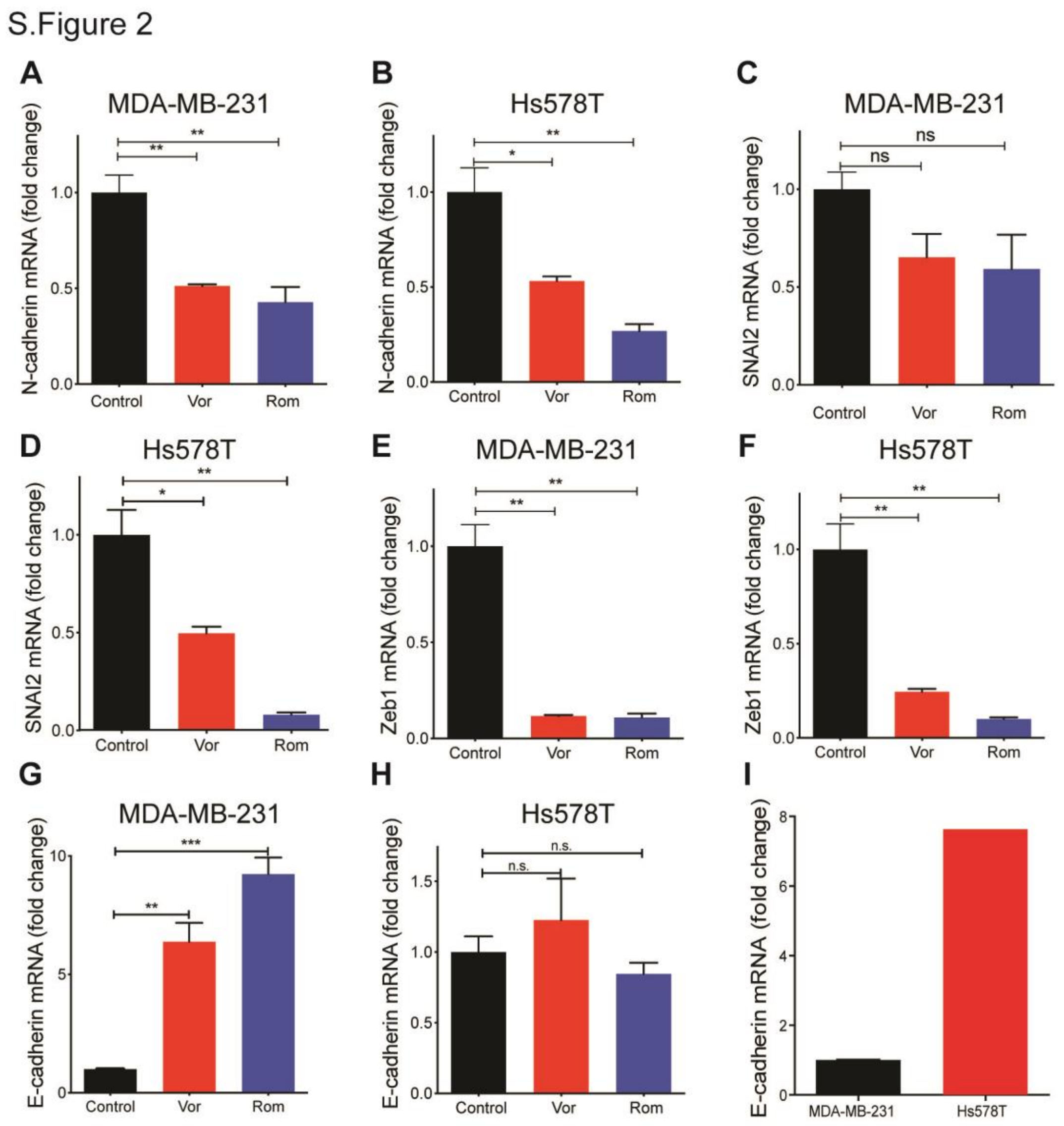


**Supplementary Figure 2: HDAC inhibitors promote MET phenotype TNBC cells**

(A-F) Treatment of MDA-MB-231 and Hs578T cells, as shown, with vorinostat (2.5 μM) and romidepsin (5 nM) for 24 hours decreases expression of several mesenchymal gene markers. Transcripts were determined by qPCR, n=3. (G-H) Treatment with vorinostat (2.5 μM) and romidepsin (5 nM) for 24 hours significantly increased transcript of the epithelial gene marker E-cadherin (CDH1) in MDA-MB-231 cells, but not in Hs578T cells; n=3. (I) Hs578T has 7-fold higher endogenous CDH1 expression compared to MDA-MB-231, in the absence of HDACi. Student t-test, **$p<0.01$,*$p<0.05$, $^{ns}p>0.05$.

S.Figure 3

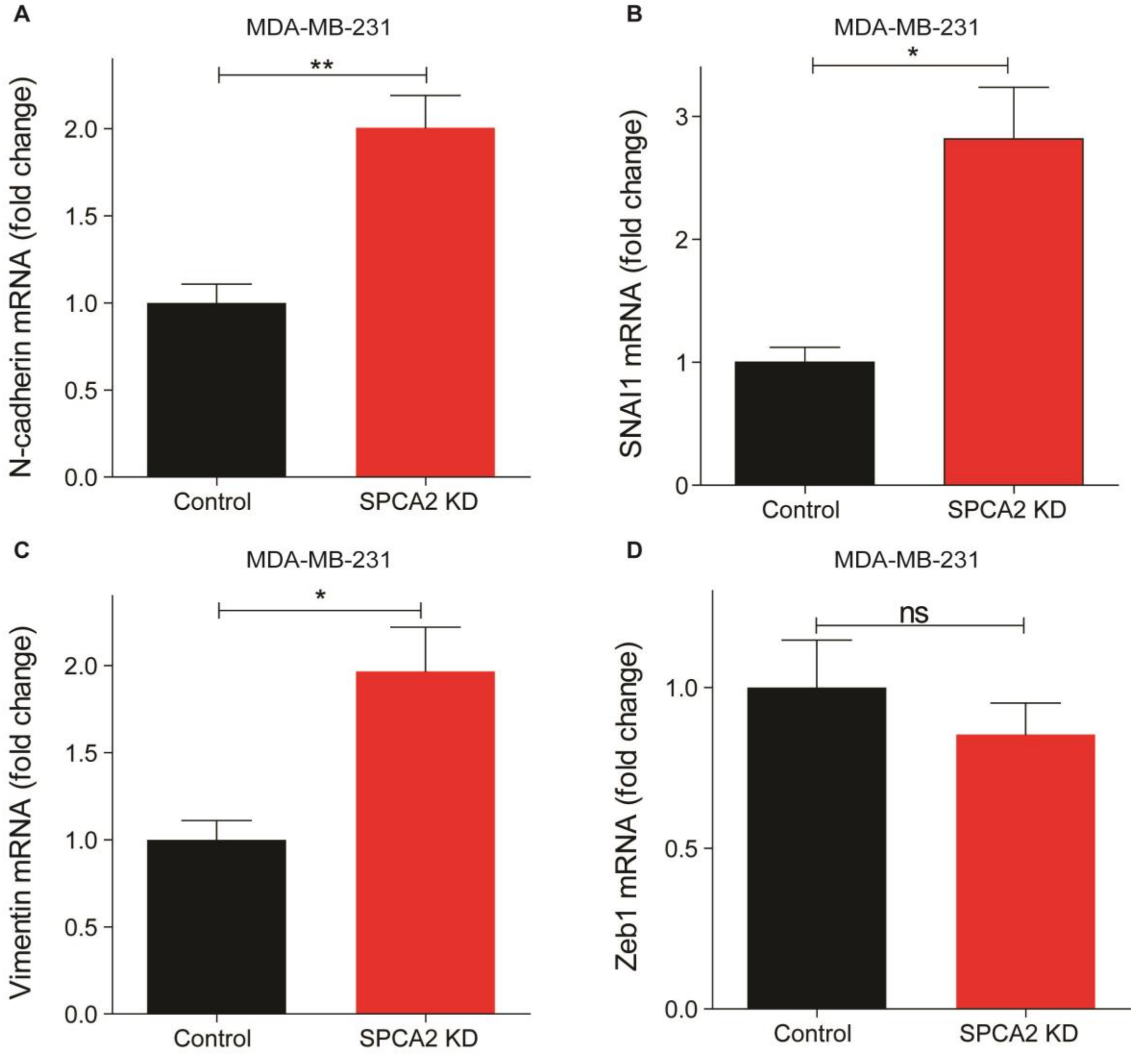


**Supplementary Figure 3: SPCA2 KD induced EMT changes in TNBC cell lines**
(A-C) Knockdown of SPCA2 in MDA-MB-231 increased mesenchymal gene markers N-cadherin, SNAI1 and vimentin. (D) Knockdown of SPCA2 in MDA-MB-231 did not significantly increase mesenchymal gene marker Zeb1. n=3. **$p<0.01$,*$p<0.05$, $^{ns}p>0.05$.

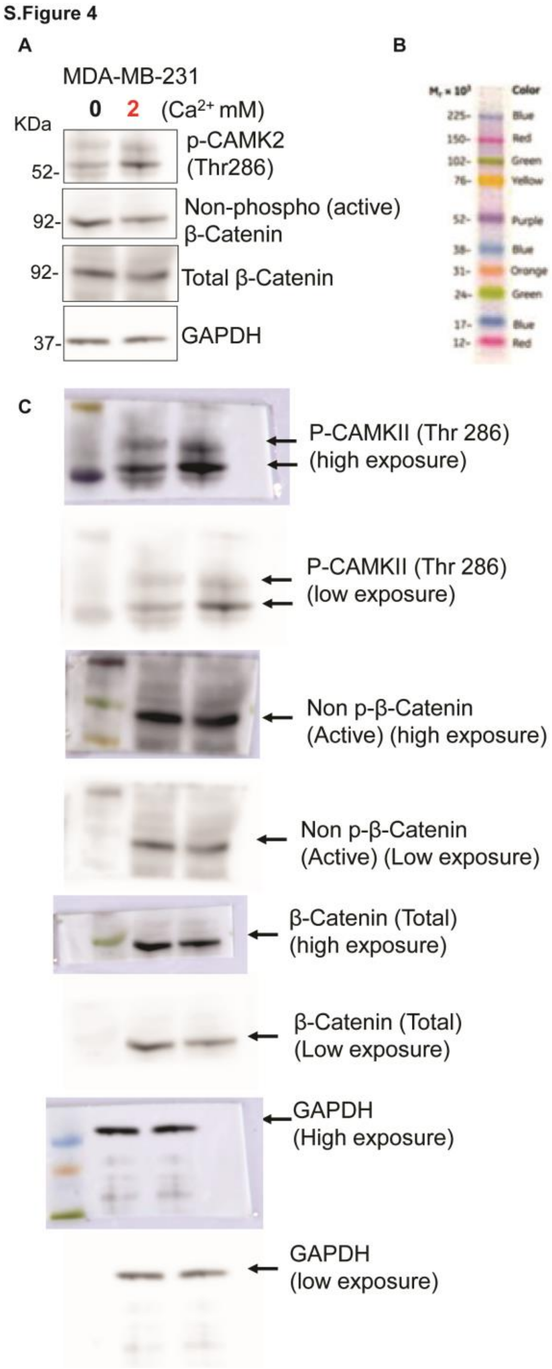


**Supplementary Figure 4: Extracellular Ca entry can also activate non-canonical Wnt/Ca pathway.**

(A) MDA-MD-231 were maintained in DMEM with 0 mM and 2 mM calcium for 42 hours. Extracellular Ca entry activated non-canonical Wnt/Ca pathway. (B) Amersham™ ECL™ Rainbow™ Marker (RPN800E, Sigma Aldrich) was used for western blots. The figure shows full molecular weight range. (C) These blots complement the cropped blots shown in Supplement Figure 4A.

S.Figure 5

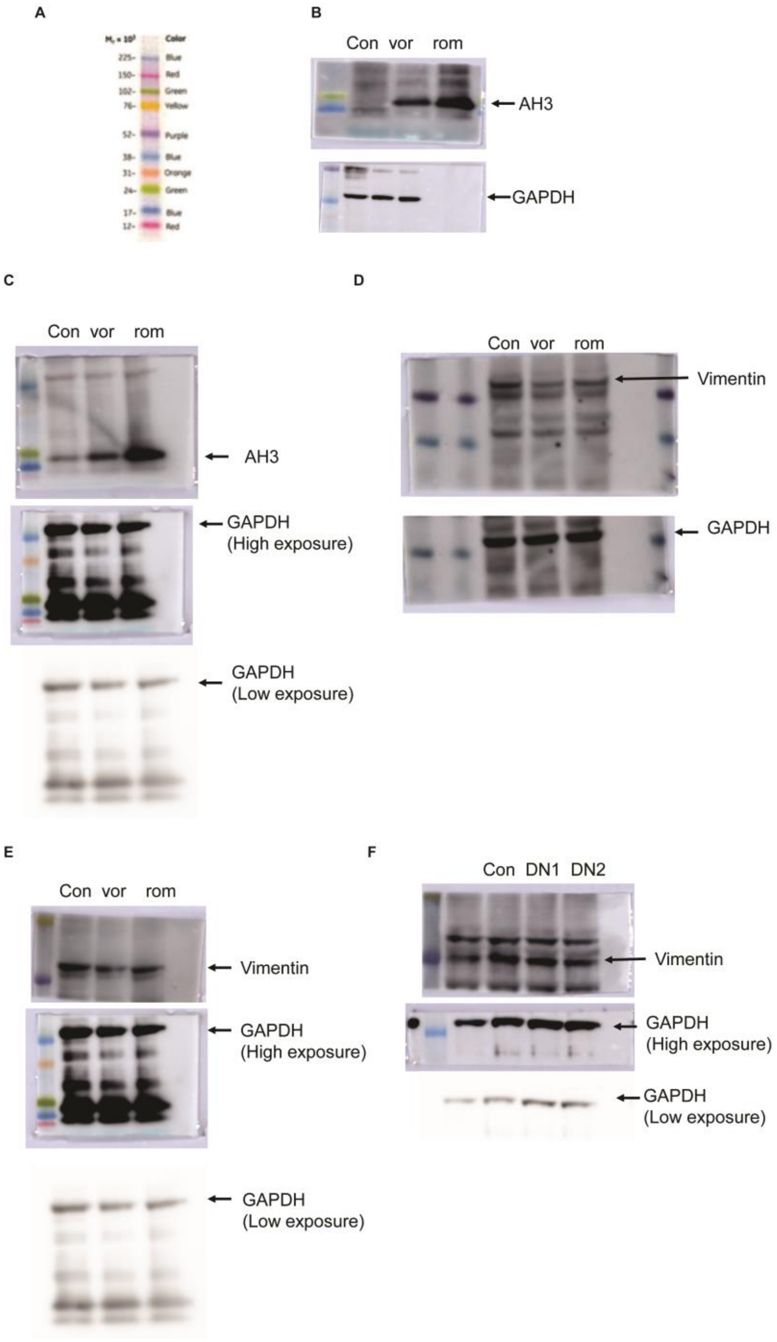


**Supplementary Figure 5: Uncropped western blots for Figures 2, 3 and 5**

(A) Amersham™ ECL™ Rainbow™ Marker (RPN800E, Sigma Aldrich) was used for western blots. (B) These blots complement the cropped blots shown in Figure 2B. (C) These blots complement the cropped blots shown in Figure 2B. (D) These blots complement the cropped blots shown in Figure 3D. (E) These blots complement the cropped blots shown in Figure 3D. This blot was developed using same sample as Supplement. Figure 5C, so loading control is same as Supplement. Figure 5C. (F) These blots complement the cropped blots shown in Figure 5B.

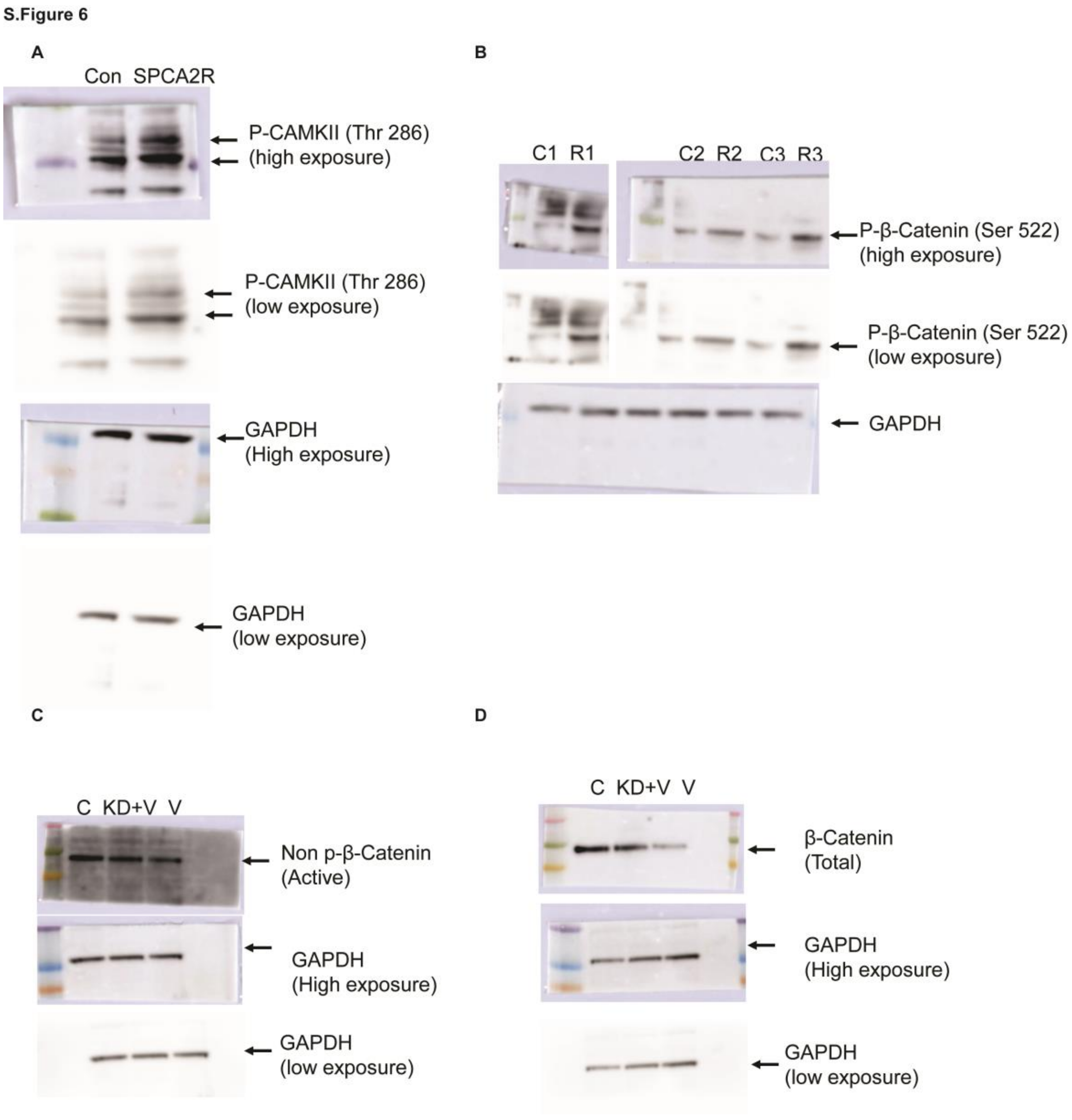


**Supplementary Figure 6: Uncropped western blots for Figure 6**
(A) These blots complement the cropped blots shown in Figure 6B. (B) These blots complement the cropped blots (C3 and R3) shown in Figure 6B. (C) These blots complement the cropped blots shown in Figure 6J. (D) These blots complement the cropped blots shown in Figure 6J.
